# Genetic load and mutational meltdown in cancer cell populations

**DOI:** 10.1101/193482

**Authors:** Yuezheng Zhang, Yawei Li, Xu Shen, Tianqi Zhu, Yong Tao, Tao Li, Xueying Li, Di Wang, Qin Ma, Zheng Hu, Jialin Liu, Caihong Zheng, Jue Ruan, Jun Cai, Chung-I Wu, Hurng-Yi Wang, Xuemei Lu

**Author notes:** These authors contributed equally to this work (Yuezheng Zhang, Yawei Li, Xu Shen, Tianqi Zhu). (HYW); (XL).

## Abstract

Large and non-recombining genomes are prone to accumulating deleterious mutations faster than natural selection can purge (Muller’s ratchet). A possible consequence would then be the extinction of small populations. Relative to most single-cell organisms, cancer cells, with large and non-recombining genomes, could be particularly susceptible to such “mutational meltdown”. Curiously, deleterious mutations in cancer cells are rarely noticed despite the strong signals in cancer genome sequences. Here, by monitoring single-cell clones from HeLa cell lines, we characterize deleterious mutations that retard cell proliferation. The main mutational events are copy number variations (CNVs), which happen at an extraordinarily high rate of 0.29 events per cell division. The average fitness reduction, estimated to be 18% per mutation, is also very high. HeLa cell populations therefore have very substantial genetic load and, at this level, natural population would likely experience mutational meltdown. We suspect that HeLa cell populations may avoid extinction only after the population size becomes large. Because CNVs are common in most cell lines and cancer tissues, the observations hint at cancer cells’ vulnerability, which could be exploited by therapeutic strategies.

## INTRODUCTION

Cancer initiation requires tumor cells to obtain several key traits, such as sustained proliferative signaling, resistance to cell death, and accelerated cell growth leading to a competitive advantage over slow-growing neighboring cells (1, 2). Accelerated proliferation of cancer cells inevitably increases their underlying rate of mutations (2, 3), copy number variations (CNVs) that affect a larger fraction of cancer genomes (4-7). It has been shown that positive selected CNVs (drivers) have critical roles in activating oncogenes and in inactivating tumor suppressors (6, 8-14). While many other mutations, including CNVs, may be fitness-neutral and are termed passenger mutations (15, 16), there in fact exist a larger class of mutations that reduce cancer cell proliferation and slow cell growth (17, 18). This class of deleterious mutations has been referred to as “negative drivers” (18).

Deleterious mutations cannot be efficiently purged by natural selection in populations with no recombination. The phenomenon is referred to as Muller’s ratchet, a moving mechanism that is used to indicate the irreversible accumulation of deleterious mutations (19, 20). Since recombination, which can enhance the efficacy of selection against deleterious mutations (20, 21), is absent in the clonally-reproducing cancer cells, accumulation of deleterious mutations can collectively exert a noticeable effect on fitness (22). Deleterious mutations may have a variety of effects on tumorigenesis by shaping cancer growth and intra-tumor variation. If the incidence of deleterious mutations is low or negligible, cancer progression can be easily described by relatively simple mathematical models and genetic variation within tumors can be treated as a function of population size. At another extreme, if deleterious mutations are prevalent, the majority of cancer cells would be defect with limited proliferative ability and only a small fraction of cells in a tumor would be capable of constantly dividing. Accordingly, the proportion and the effect size distributions of deleterious mutations must not be overlooked and may be key parameters to model tumorigenesis.

Guided by previous experience in population genetics, most studies in cancer focus on slightly deleterious mutations as they should be able to evade purifying selection and accumulate, thus influencing long-term tumor progression (23). Because highly deleterious mutations should be rapidly weeded out by purifying selection, it is generally assumed that these mutations have little to do with long-term cancer progression. Thus, the most common mutations, either fixed or at high frequency, that are frequently cited in cancer genomic studies are usually advantageous for individual cell fitness. Such practical limitations ensure that deleterious mutations are mostly undetected in empirical studies. Indeed, even the relatively comprehensive TCGA pan-cancer data set yielded only a small number of deleterious mutations, highlighting gaps in our knowledge about this important class of variants (18).

Direct measurements of individual deleterious mutation effects *in vivo* are challenging. However, an assessment of their collective action is possible, given a system that generates such variants at an appreciable rate. HeLa cells present such as system as their genomes are highly variable. In addition, this cell line has attractive features for our purposes since it has been extensively cultured and exhibits a short doubling time and aggressive growth. We examined variation in growth rate among individual HeLa cells by monitoring clones from a common ancestral HeLa cell population. We then estimated deleterious mutation rate and the average fitness decrease per mutation by performing computer simulations of cell growth. Our observations suggest that the differences in growth rate among cell clones are heritable, and CNV is the major cause of proliferative fitness reduction in HeLa cells. HeLa cells constantly produce high numbers of deleterious mutations during growth. We discuss the implications of our findings for modeling tumorigenesis.

## RESULTS

### Genetic variation in growth rate in a HeLa cell population

To ensure genomic homogeneity of the initial population, we first established a HeLa cell line (E6) derived from an ancestral cell line (JF) (Fig. S1). When E6 population size reached approximately 5 × 10^4^ cells (15∼16 divisions), five single-cell clones were generated from E6 and established in culture. When the clone cultures reached 10^6^ ∼ 10^7^ cells, we measured their growth rates using the MTT assay. The B8 and G3 clones showed clearly higher growth rates than E3, E7 and G2, suggesting that cells within E6 were heterogeneous (Fig. 1A). Since these clones were all descendants of E6 which originated from a single cell and experienced only 15 - 16 divisions, our results indicate that heterogeneity in growth rate can be generated in a very short period of time in cancer cells.

**Fig 1.**
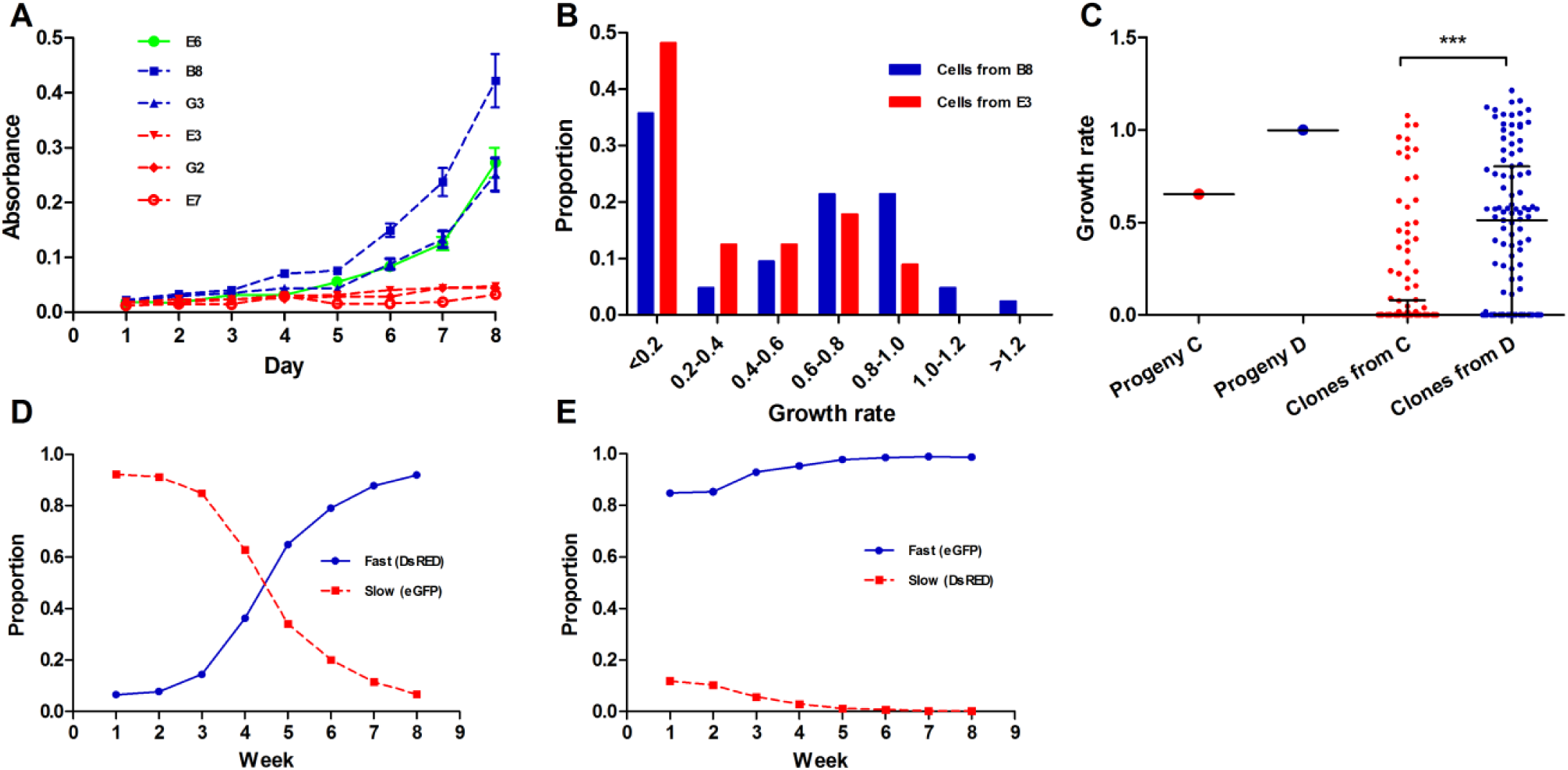
Growth rate, survival rate and fitness of fast- and slow-growing cells. (A) Growth graphs of 5 descendant clones (B8, G3, E3, E7, G2) and the ancestral clone (E6). The mean MTT assay read-out (the values of absorbance, y-axis) taken from day 1 to day 8 were plotted for each clone (8-12 replicates). Blue red and green lines represent growth graphs of fast, slow and ancestral clones, respectively. Error bars represent standard errors. (B) Proportion of progeny with different growth rates. The growth rates of 40 B8 (red) and 39 E3 (blue) single cell progeny were monitored and calculated for seven days. (C) Growth rate of one slow-(C; red) and one fast-(D; blue) growing descendant B8 and its single-cell clones. Average growth rate of the cell clones (blue dots) from D is significantly higher than that of the clones (red dots) from C (p = 2 × 10^−11^, KS test). (D and E) Competition assay between slow- and fast-growing cells. The proportion (y-axis) of fast-(blue) and slow-growing (red) cells in a mixed population was measured from week 1 to week 8 (x-axis) by flow cytometry.

To test whether variation in growth rate among clones is heritable, we isolated 39 cells from B8 (fast growing clone) and 40 cells from E3 (slow growing clone), and monitored their growth from a single cell for seven days. Approximately 30% of B8 and 50% of E3 cells died out within seven days (Fig. S2), due to either damage caused during cell isolation or genetic defects. Furthermore, most cell lines with growth rates < 0.6 died within 2 months. Thus, only 50% of B8 and 27% of E3 cells survived for more than two months.

While the growth rates of cells from a single origin varied greatly, the progeny of the fast-growing clone (B8) grew faster on average than those of the slow growing clone (E3)(Fig. 1B). Mean cell number among the B8 progeny was 62.0, while it was 17.3 among E3-derived cell lines. We further drew approximately 100 cells from one fast-(progeny D) and one slow-(progeny C) (Fig. S2) growing derivative of B8 and monitored growth of each cell clone for seven days. As expected, cells from the fast-growing progeny had a higher mean growth rate than that from the slow-growing descendant (*p* = 2 x 10^−11^, KS test; Fig. 1C).

If mutations that slow cell proliferation frequently arise in cancerous cell populations, we would expect a decrease in proliferation rates and an increase in among-cell growth speed variance as the population is maintained. To test this, we monitored growth of single cells that were randomly drawn from cell populations at different time points. We first set up six single-cell clones from B8 (Table S1). After cell numbers reached 100 - 500 (8-10 cell divisions), 20 - 30 % of the cells from each population were randomly drawn and separated into single cells. The growth of these isolates was monitored for eight days. This step was repeated when the size of the six populations exceeded 5,000 cells. In all six cases, the average growth rate of cells drawn at the first time point was higher than those from the second time point (Fig. S3). In addition, in four out of six cases, variation in growth rate was higher at the second than at the first time point (*t*-test).

To test whether the slow-growing cells would be outcompeted by their fast-growing counterparts, we performed a competition assay. The fast- and slow-dividing cells were co-cultured in different ratios and their proportions in populations were monitored weekly using the *Discosoma* sp. red fluorescence protein (DsRed) and enhanced green fluorescence protein (eGFP) over time using flow cytometry. We ran the experiment for eight weeks. Regardless of the initial proportions, the fast-dividing cells always outgrew the slow-dividing cells in our co-culture assays over time (Fig. 1D and E), suggesting that fast cells indeed possessed higher fitness than the slow ones.

Fig. 1 demonstrates that a cancer cell population can generate heterogeneity in growth rate within several cell replications, even starting from a single cell. Moreover, the majority of these changes is heritable and reduces the fitness of cancer cells (defined as proliferation rate), suggesting that fitness reduction we observe is largely genetically determined.

### Cell growth rate is associated with CNV number

To study the genetic basis of growth rate heterogeneity among our cancer cell lines, we assessed genomic variation in E6 and five of its descendant clones by constructing a digital copy-number profile based on next generation sequencing reads. We focused on copy number variation because single-nucleotide mutation rates are too slow to produce significant sequence variation during our short-duration culturing experiments. Most clone genomes are triploid (Fig. S4), and only a small number harbors more than four or fewer than two haploid genomes, consistent with a previous study (24).

The three slow-growing clones showed a clear increase in CNV copy number compared to the E6 parental line (Fig. 2, Table S2), while the two fast-growing clones were more similar to E6, suggesting that that most CNVs are deleterious. As E6 experienced only 15 - 16 divisions before its five descendant clones were generated, above results also indicate that CNVs can be generated in a very short period of time. To test this hypothesis, 11 clones derived from B8 with different growth rates between day 1 to 8 were picked for further analyses (Fig. 3A). The growth rate of each clone was measured again by RTCA iCELLigence when the population reached approximately 10^6^ cells (after 20 - 30 cell divisions). The results were highly correlated (R^2^ = 0.713, p-value < 0.01; Fig. 3B) with the previous eight day measurements, demonstrating that variation in growth rates among clones was consistent and not due to stochastic fluctuation at different stages.

**Fig 2.**
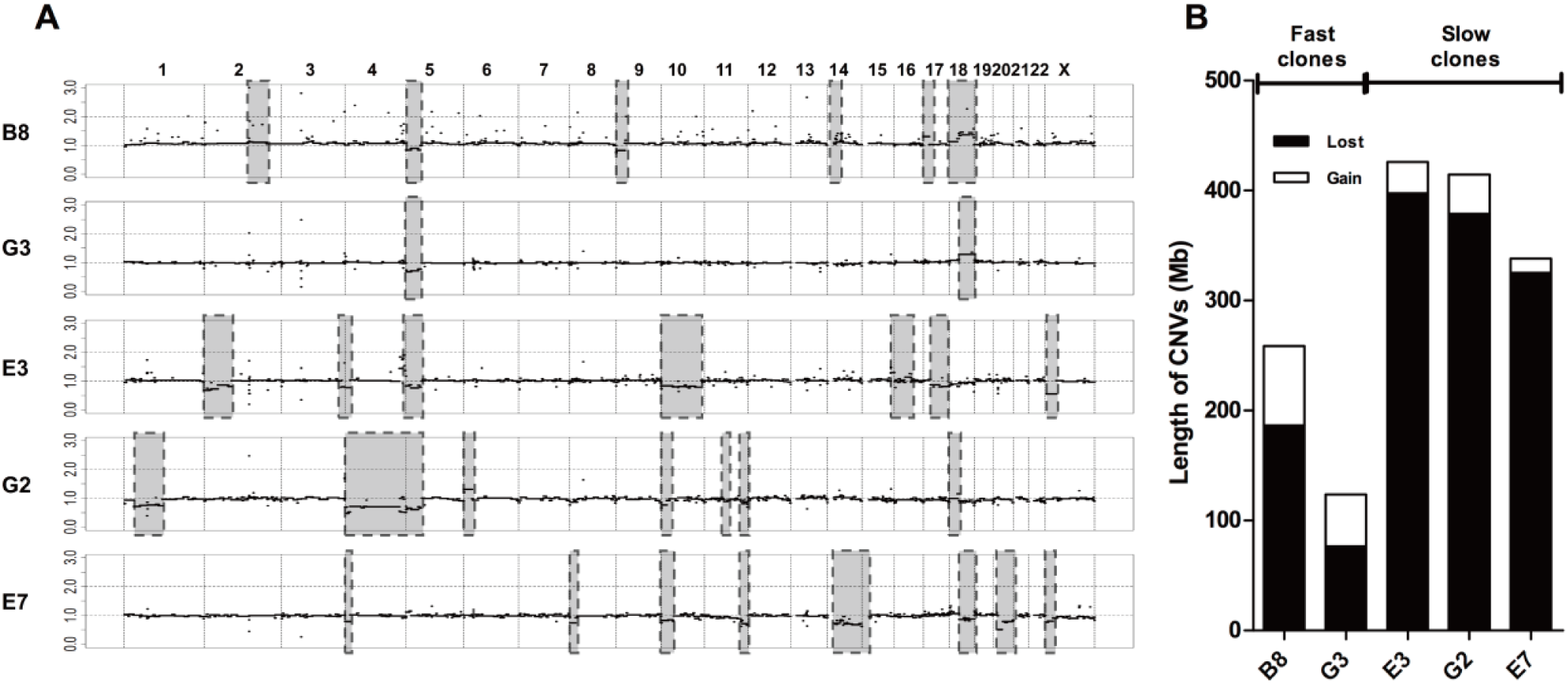
Copy number variation (CNVs) in five clones derived from E6. (A) The CNV regions in fast-(B8 and G3) and slow-growing (E3, E7 and G2) clones are highlighted with grey rectangles. The y-axis is the ratio of copy number in a descendant clone and copy number in the ancestor, E6. (B) Summary of CNV gain and loss among five descendant clones (see also Table S2).

**Fig 3.**
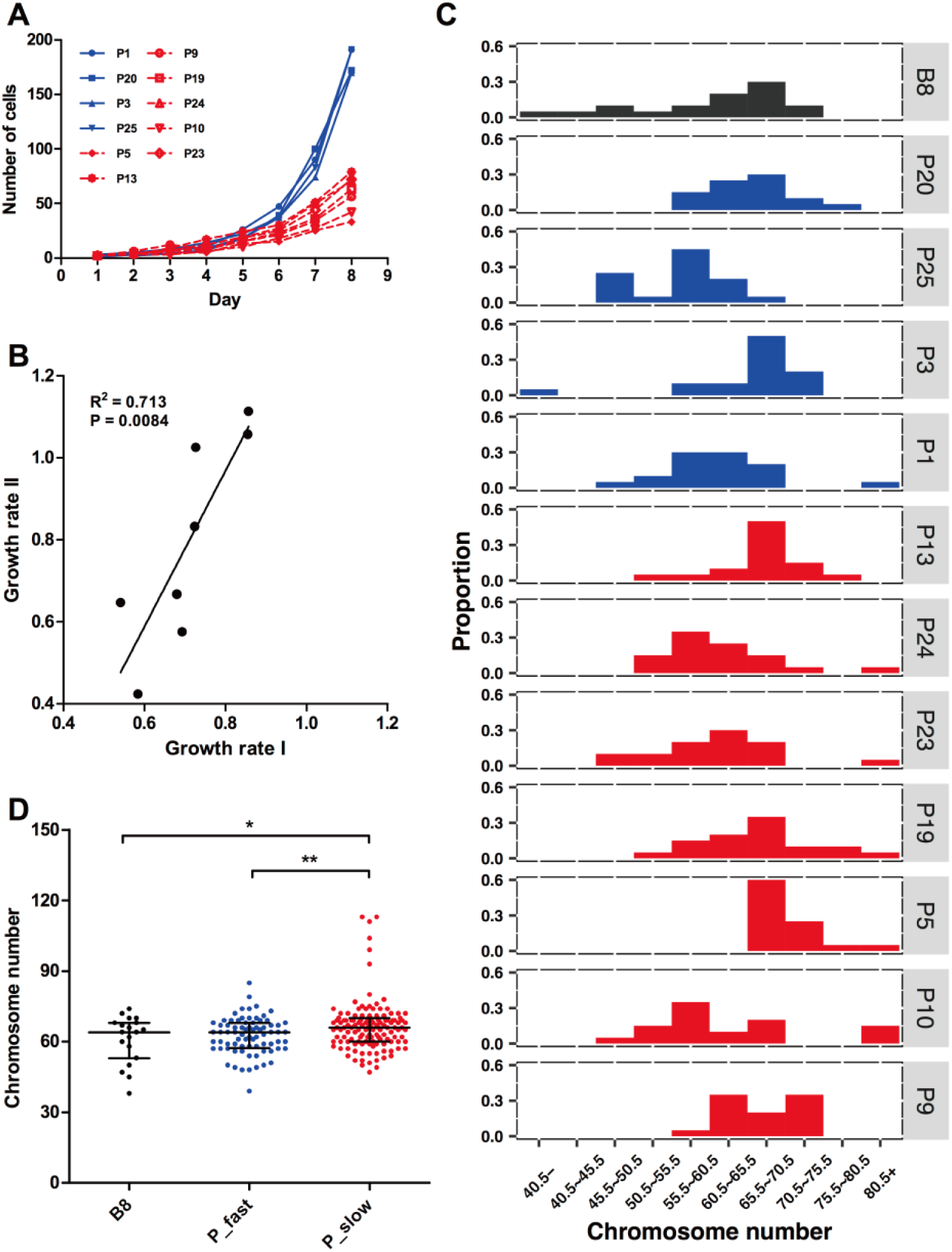
Growth rate and chromosome number variation among single-cell clones generated from B8. (A)Cell numbers of single cell clones from day 1 to day 8. The clones labeled in blue grow faster than the clones in red. (B) Correlation between growth rate I and II of single cell clones. The growth rate I (x-axis) was measured in the first eight days and the rate II (y-axis) measured by RTCA iCELLigence method when the cell populations reached ∼10^6^ cells. Each dot represents a single cell clone; only eight out of 11 clones were measured at the second time point. (C) Distributions of chromosome numbers in cells from ancestal and descendant clones. Chromosome numbers (x-axis) of 20-30 cells randomly drawn from each clone were counted. The black, blue, and red histograms represent cells from the ancestral, fast, and slow progeny clones. (D) Chromosome number in cells from the ancestral (B8) and the progeny clones. The black, blue, and red dots represent cells from the ancestral, fast (P_fast), and slow (P_slow) progeny clones. *: p < 0.05, ** p < 0.01.

We picked about 20 cells from each of the 11 clones and counted their chromosome numbers. The karyotypes ranged from 38 to 113 chromosomes, with most (72%) cells harboring between 55 and 70 chromosomes (Fig. 3C). Therefore, despite single-cell origin, the progeny quickly generated aneuploidy within only 20-30 cell divisions, again illustrating frequent cytogenetic change in cancer cells.

Although chromosome numbers varied among clones, their distribution by combing all clones reconstituted the chromosome distribution of their ancestor B8. The average number of chromosomes in the fast-growing group (62.5) was similar (p = 0.56, t test) to the B8 clone (61.1), whereas the slow growing group (66.5) showed significantly greater chromosome numbers than the fast growing (p < 0.01) and the ancestral (p = 0.04) clones (Fig. 3D). Consequently, Fig. 2 and 3 suggest that cancer cells exhibit a very high rate of CNV generation and most of these CNVs are deleterious, reducing the cells’ proliferative ability.

### Rate of deleterious mutations

To further study how fast the difference in proliferation rate can be generated, we picked 12 cells from B8, allowed them to divide once, and isolated each daughter cell in a well of a 96-well plate. We then monitored the growth of each of the 12 pairs of cells for four days (Fig. 4A). Four cell pairs lost at least one of the daughters in the first four days, probably due to injury during preparation. Of the remaining eight pairs, two (h and i, Fig. 4A) exhibited different proliferation rates between the daughter cell lineages. The result suggests that there is approximately one deleterious mutation in every four cell divisions in this cancer cell line.

**Fig 4.**
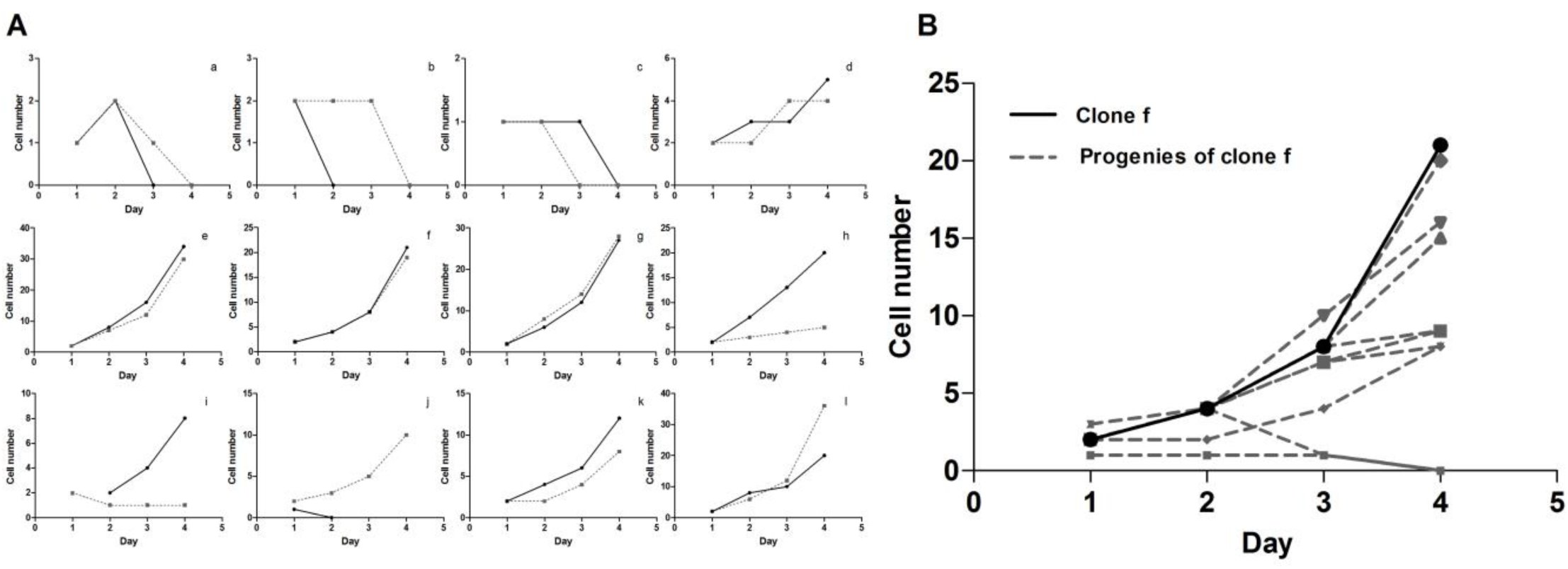
Proliferation rate differences between pairs of daughter cells. (A) Growth graph of two daughter cells (solid and dashed line) derived from a single cell. The number (y-axis) of cells in 12 pairs (a to l) of daughter cells was counted every day in four days. (B) Growth graph of progeny cells from clone f. The progeny cells were isolated from clone f and monitored for four days. Solid line: clone f; Dashed line: progenies of clone f.

Such a high deleterious mutation rate implies that many cells at day four would carry deleterious mutations which reduce cell proliferation. To test this, each descendant of clone f was harvested and re-suspended into a 96-well plate. We monitored growth of these progeny for another four days. As expected, the majority of the surviving clones exhibited slower growth than their ancestor f (Fig. 4B).

### Modeling population growth attenuation

Rough estimates from our experiments so far suggest extremely high mutation rates in cancer cell lines. To understand mutation accumulation rates in tumors, we need quantitative estimates of deleterious mutation rates and effect distributions. Therefore, we constructed and applied a simple model of cell growth and the mutation accumulation process (see *Materials and Methods*). Let *N*_*t*_ be the population size at day *t*, where *N*_*0*_ = 1, and *R*_*t*_ be the population growth rate at day *t*. For each generation, a proportion of cells (*μ*) generates new mutations which decrease their growth rate. The mean deleterious effect of a mutation is *d*. We have

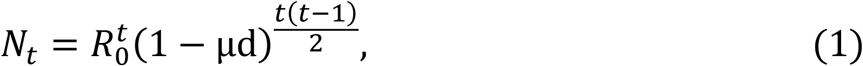

where *R*_*0*_ is the net growth rate at day 0.

To estimate parameters of this model (*R*_*0*_, *μ*, and *d*) we randomly drew 18 single cells from B8 and monitored their growth for 7-8 days (Table S3). We divided these newly-derived cell lines into fast- (cell number > 100) and slow-growing (cell number < 100) groups. We then conducted computer simulations to evaluate the parameters that best fit the observed data (see *Materials and Methods*).

Using fast-growing cell data, we estimate posterior mean of *R*_*0*_ = 2.37 ([2.22, 2.52]), *μ* = 0.29 ([0.26, 0.30]), and *d* = 0.18 ([0.17, 0.20]). Estimates from slow-growing cells (*R*_0_ = 2.00 ([0.25, 0.33]), *μ* = 0.29 ([0.16, 0.20]), and *d* = 0.18 ([2.22, 2.52])) are similar, except that the initial growth rate is slower, as expected. The deleterious mutation rate (*μ*) of 0.29 (0.26, 0.33) suggests that there is approximately one deleterious mutation for every 3 - 4 cell divisions, which is very close to our rough non-parametric estimates from daughter-cell divisions (see Fig. 4A). Since *μ* is scaled per cell division, the product of *μ* and *d* reflects fitness change per generation. Our estimates indicate that the HeLa cells experience a 5% (4%, 6%) reduction in fitness for every generation (25, 26) (Table S3). Using point estimates of *μ* (0.29) and d (0.18), we fitted our model to the growth rates observed in a range of experimental data (Fig. S2) and estimated initial growth rates (*R*_*0*_). Only cell lines that showed monotonic increase in cell numbers were considered. The estimation of *R*_*0*_, ranging from 1.64 to 2.54 in 43 sets of experimental data (Table S4), suggested that the ancestral cell of slowest growing lineage had accumulated about 2.2 (log (1.64 / 2.54) / (1 – 0.18)) more deleterious mutations than the ancestor of fastest growing lineage.

Interestingly, while *R*_*0*_ shows substantial variation among cell lines, estimates of *μ* and *d* are similar, possibly indicative of an intrinsic property of rapid cancer cell division. To further test this, we estimated the *R*_*0*_, *μ* and *d* in tumor cell lines from different cancer types by the same method (Table 1). Although the *R*_*0*_ are considerably variable, *μ* and *d* are generally consistent among cell lines, implying that a fitness reduction (*μ* × *d*) in a variety of cancer cells is close to our initial estimate of 5% per generation.

**Table 1.** Fitness reduction per cell division in different cancer cell lines.

| Cell line | Cancer type | Number of clones<sup>1</sup> | $R_0$<sup>2</sup> | μ<sup>3</sup> | $d$<sup>4</sup> | Fitness reduction ($\mu \times d$) |
| --- | --- | --- | --- | --- | --- | --- |
| Hela | Cervical cancer | 18 | 1.66 - 2.48 | 0.25 - 0.33 | 0.17 - 0.20 | 0.04 - 0.06 |
| PC3 | Prostate cancer | 10 | 1.68 - 2.11 | 0.26 - 0.31 | 0.16 - 0.20 | 0.04 - 0.06 |
| A204 | Rhabdosarcoma | 7 | 1.81 - 2.07 | 0.28 - 0.33 | 0.17 - 0.22 | 0.05 - 0.07 |
| A375 | Melanoma | 18 | 2.14 - 2.64 | 0.26 - 0.34 | 0.16 - 0.23 | 0.05 - 0.08 |

To estimate parameters in equation (1), single cells were randomly chosen from different cancer cell lines and their growth from day 0 to day 7 was monitored. Fitness reduction was evaluated based on the parameters estimated by applying computer simulations, which are described in Results and Methods.
<sup>1</sup> The number of single cell clones generated from different cancer cell lines
<sup>2</sup> Growth rate of cell population at day 0
<sup>3</sup> Deleterious mutation rate
<sup>4</sup> Mean deleterious effect of a mutation

## Discussion

We performed multiple single-cell progeny assays using HeLa cell lines and extensively genotyped the clonal populations, focusing on copy number variants (CNVs). We fitted a growth model to these data to estimate distributions of mutation effects on fitness as measured by proliferative capacity. We show that CNV deleterious mutations appear at the rate that falls in the range of 0.26 - 0.33 per genome per generation. Our results demonstrate that accumulated passengers are deleterious to cancer cells because they reduce cell proliferation. We estimate that the deleterious effect of 0.18 ([0.175, 0.187], 95% confidence interval) is twice higher than previous results (17, 27). While at first this may seem to be unreasonably high, the mutations we identified would not be seen in cancer cell populations within tumors as the vast majority of these variants are too deleterious to appear in the earlier data sets.

Nevertheless, the observation that a significant portion of cancerous cells are born with a low enough proliferative capacity that they are rapidly lost during tumor growth suggests that there is a trade-off between cancer progression and genome instability. While cancer cells may benefit from generating genetic variation paving the way for tumor progression, accumulation of deleterious mutations would eventually be intolerable and lead to cell population meltdown. The tumor population must thus strike a delicate balance to maintain heterogeneity and at the same time curb relentlessly increasing passenger mutations. Exploiting this fact may help develop a new therapeutic regime for cancer (17, 23). This can be done by either increasing the overall mutation rate (*μ*) or the deleterious effect of passengers (*d*). For example, a 10% increase in *μ* may result in a more than 50% population reduction in 20 days (Fig. S5). Practically, mutation rate can be increased by targeting the DNA repair system (28) or by regulating DNA replication stress (29).

However, for these strategies to be effective, an additional layer of cancer biology needs to be considered. It has been shown by mathematical modeling that a high incidence of defective cells and cell death may not be disadvantageous for tumor growth, but in fact facilitate tumor progression (30). That is because a high rate of proliferation-reducing or cell death-inducing spontaneous mutation can lead to cancer dynamics that enable sufficient stem-like cell divisions to enrich the pool of cells with high proliferation rate and drive population expansion.

Therapies that directly target cell proliferation should be considered. Since in order to maintain high proliferation rates, rapidly proliferating cells need to increase their translational capacity and are dependent on high rates of ribosome biogenesis (31, 32). Thus, inhibition of ribosome biogenesis could be a selective approach to cancer therapy (33, 34). More importantly, this effect is enhanced in cells with higher proliferation rates, but less so in cells with lower proliferation rates.

## Materials and Methods

### Cell culture and Single cell isolation

Cancer Cell lines, including HeLa cell lines, Human prostate cancer cell line PC3, Human Rhabdomyosarcoma cell line A204 and Huma malignant melanoma cell line A375, were obtained from the Culture Collection of the Chinese Academy of Sciences, Shanghai, China. Cells were cultured in DMEM Medium for HeLa cell, PC3 and A375, in medium of McCoy’s 5A for A204 (Life Technologies) and supplemented with 10% fetal bovine serum (FBS) (Gibico, Life Technologies), 100 U/ml of penicillin, and 100 μg/ml of streptomycin at 37°C with 5% CO_2_. Cells were trypsinized using 0.05% trypsin at room temperature for 3 min. Cell counting was performed with a hemocytometer. The cell suspensions were diluted with medium to the final concentration of 1 cell per 100 μl. Single cells were seeded into each well of a 96-well cell culture plate and maintained at 37°C with 5% CO_2_. After incubating for 12 hours, during which the cells attached to the wells, the single cell isolation was visually confirmed by microscopy. The number of cells in each well was counted every day. When the cell colony became sufficiently large, cells were transferred to 24-well plates, and subsequently transferred to 6-well plates and 10-cm culture dishes.

### Karyotype analysis

The cultured cells in a stage of active division were treated with colchicine (200 ng/ mL) for 1 hour at 37°C, then harvested and resuspended in 0.07M KC1 for 30min, and slowly added 10 drops of Carnoy’s fixative (3:1, methanol:acetic acid). The suspended cell was dropped onto a slide, dried the slide rapidly, and stained with 4% DAPI for 5 min. At least twenty cells were spread out in metaphase for karyotyping. Evaluation of interphase nuclei was performed by OLYMPUS BX51 fluorescence microscope. Photographs were taken by a CCD camera with 40 or 100 times objective. Image-Pro Plus software was used for digital image acquisition in the TIFF format, pseudocoloring and merging.

### MTT Assay

MTT assay was used to measure cell proliferation. Cells were suspended and seeded at the concentration of ∼700 cells/100μl/well in 96 well plate. A volume of 20 μl dissolved MTT was pipetted into each well. After being incubated for 4 hours at 37°C in a humidified CO_2_ incubator, the media was removed and 200 μl sterile DMSO was added to each well. The absorbance values were then read at 570 nm with a microplate spectrophotometer. The number of living cells was estimated based on absorbance values.

### Real-time cell analysis assay (RTCA)

Cells in 10% FBS media were seeded at a density of 5000 per well into a 16-well E-plate (ACEA) and incubated for 72 hours at 37 °C in a humidified CO2 incubator. The impedance for each well was monitored by the RTCA iCELLgence™ system (ACEA Biosciences Inc) every 70 minutes. Relative impedance signal level (represented as “cell index” in manufacturer’s software) that indicated the number of cells was analyzed using the RTCA Software 1.2 program (Roche Diagnostics). The cell growth curves were automatically recorded on the iCELLigence system in real time. The cell index was followed for 3 days.

### Measurement of growth rate

The single cell clones that we generated for the growth rate calculation were in the log phase, the period of exponential increase in the cell number. During this phase,

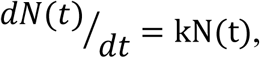

where N(t) is the total number or the concentration of cells at time t, and k is a constant coefficient. To obtain a linear function, the cell numbers were converted to logarithms to the base of Euler’s number e. The least-squares method (LSM) was used to fit the linear regression in which the slope (k) of the regression line would be the growth rate.

### Whole genome sequencing and analysis of copy number variation

Genomic DNA was extracted from 10^6^ HeLa cells using QiagenAllpre DNA/RNA Mini Kit (Qiagen). The genomic DNA (3 μg) was fragmented by Adaptive Focused Acoustics on a Covaris E120 (Covaris Inc). The range of product size was from 250bp to 350bp. The fragmented DNA was purified by Qiaquick PCR purification column and quantified on 2100 Bioanalyzer by using the Agilent DNA 1000 kit (Agilent Technologies, Palo Alto, CA, USA). The whole genome libraries were constructed by IlluminaTruseq DNA sample preparation kit according to the manufacturer’s instructions. Whole genome sequencing was performed using the Illumina Hiseq 2000 in the Beijing Institute of Genomics (BIG, Beijing, China). Reads were mapped to the human reference genome (hg18) using BWA software (Version 0.4.9) with default parameters (35). The aligned reads were used as input of the Control-FREEC software (Control-FREEC v10.3; http://bioinfo-out.curie.fr/projects/freec/) for characterizing the large-scale chromosomal and segmental duplication and deletion events (> 10^7^ bp) in all those samples (Fig S4). FREEC was run with the following parameters: window size, 100kb; step size, 50 kb; contaminationAdjustment = TRUE; noisyData = TRUE; BAF calculation inactivated. The segmental copy number gain and lose of each descendant clone was profiled against the E6 genome by preforming the FREEC with the same parameters mentioned above. The relative ratios and breakpoints of copy number gain and lose against the E6 were calculated and identified for the 5 descendant clones (Fig 2A, Table S2).

### eGFP and DsRed transfection

Cells with the stable expression of eGFP and DsRed were constructed by transfecting with lentivirus vectors, plenti6.3-MCS-IRES-eGFP and plenti6.3-MCS-IRES-DsRed (Life Technologies), which contained a blasticidin resistance gene and an enhanced green fluorescent protein (eGFP) or a discosoma sp. red fluorescent protein (DsRed) sequence. The expression of eGFP and DsRed were used for labeling and distinguishing the two cells in the competition assay. The vectors were packaged into the lentivirus particles with infectious activity (Invitrogen). Before transfection, 2×10^5^ cells per well were incubated with DMEM in a 6-well plate. After incubating for 24 hours, the medium was replaced by the transfection medium that contained the lentivirus particles and polybrene with the concentration of 8 μg/ml. The multiplicity of infection (MOI) value was 3. After transfecting for 24 hours, the cells were washed three times with PBS, and cultured in the DMEM medium with blasticidin of 10 μg/ml for at least 4 weeks in order to select cells that stably express eGFP and DsRed.

### Isolation of fast dividing and slow dividing cells

The CellTrace^™^ Violet Cell Proliferation Kit, for flow cytometry (Thermo Fisher, C34557) was used for isolating the fast growing and slow growing cells. Cells were labeled by the cell-dye following the CellTrace™ Violet Cell Proliferation Kit workflow after cell cycle synchronization arrested at G1/S phase (double thymidine block). The labeled cells were continuously cultured for 7 days. At the 7th day after labeling, the cells were detached by trypsin-EDTA solution and suspended on culture medium. The BD Influx flow cytometer (BD) was used to isolate the fast growing and slow growing cells. The top 10% cells with strong fluorescence signals were sorted as the slow growing cells and bottom 10% cells with weak fluorescence signal were sorted as the fast growing cells. Analysis was completed using the BD Influx flow cytometer with 405 nm excitation and a 450/40 nm bandpass emission filter.

### Competition assay for slow growing cell and fast growing cells

Fast-dividing cells with stable eGFP expression (or DsRed expression) and the slow-dividing cells with DsRed expression (or eGFP expression) were mixed and co-cultured at different initiation ratios (1:1, 2:8 and 8:2). The proportions of the two cell populations in the mixture were monitored by fluorescent intensity of DsRed and eGFP every 3 days by using flow cytometry.

### Modeling population dynamic of cell expansion

We construct and apply a simple model of cell growth and mutation accumulation process. We define *N*_*t*_ as population size at the day t, where *N*_0_ = 1, and *R*_*t*_ as the growth rate of population at day *t*. In this case, *R*_*t*_ > 1 represents population increase and *R*_*t*_ < 1 means population decrease. For each division, a proportion of cells (μ) generate new mutations which decrease growth rate of the cells. The deleterious effect of a mutation is *d*. *N*_*t*_ is denoted by

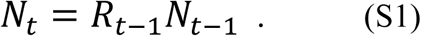

*R*_*t*_ and *R*_*t*-1_ are given by a recursive function

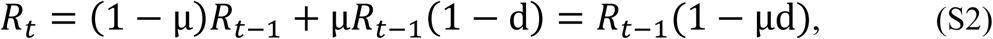

Iterating (S2) t times, *R*_*t*_ can be expressed as

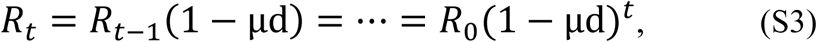

where *R*_*0*_ is the growth rate at day 0. Substitute (S3) into (S1) yields:

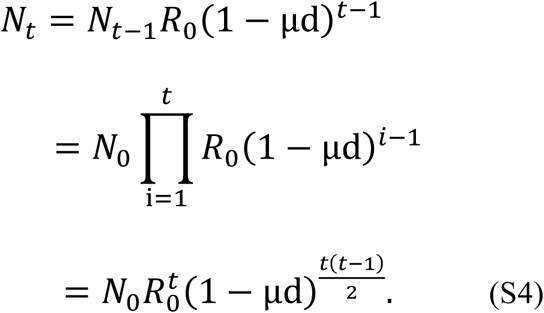

Since the experiments start from single cell isolation (*N*_*o*_ = 1), *N*_*t*_ can be derived as

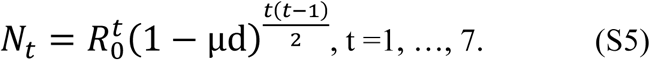

### Estimate *R*_0_ and fitness reduction μd

In (S5) *N*_*t*_ (t = 1,…, 7) can be obtained from observed data, so *R*_0_ and *μd* can be expressed by *N*_*t*_ (t = 1,…, 7). We then divide *N*_*t*-1_ by *N*_*t*_ and divide *N*_*t*_ by N_*t*+1_

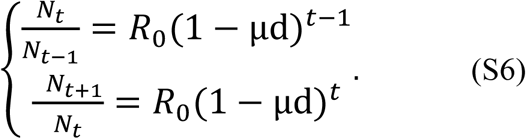

Combine two formulas in (S6) and eliminate *R*_0_, the fitness reduction μd can be calculated as

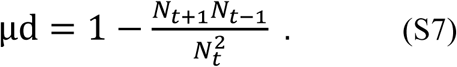

Using linear regression model with the cell number from 8 days, the fitness duction μd can be approximated as

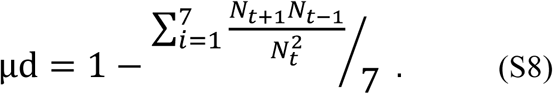

To calculate *R*_0_, we substitute (S7) into (S6). Then *R*_0_ can be expressed as

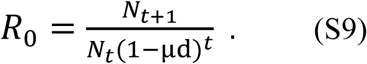

### Cell growth simulation

We extended the previously described genetic load to simulate the cell growth with accumulation of deleterious mutations. The simulation begins with a single cell. Each day, cells attempt to divide and produce additional cells. The initial growth rate, *R*_*0*_ (*R*_0_ ϵ [1.0,3.5]), follows a Poisson distribution. For each cell replication at day *t*, the number of growth rate (*R*_*t*_) which is randomly generated followed by mean growth rate *R*_*t*_. *R*_*t*_ > 1 represents cell proliferation, while *R*_*t*_ = 0 means cell death. During replication, a proportion of cells (*μ*) generates deleterious mutations. The mutation accumulation in cells leads to the decrease in growth rate. The average deleterious effect of a mutation is *d*. Therefore, the growth rate of a cell population is reducing as a function of time. The computational model is qualitatively similar to our mathematical model, but considers much more complicated conditions, i.e., the fluctuations of the parameters (*R*_*t*_, *μ* and *d*) caused by their distributions.

When fixed *R*_0_, *μ* and *d* are given, we can monitor the cell number every day using the simulation model. In summary, we cultured 10^6^ cell lines in the simulation with different combinations of *R*_0_, *μ* and *d*. Compared the 10^6^ results with observed data with HeLa cell lines, we used the Approximate Bayesian Computation (ABC) method to select optimal groups. The mean number of the selected groups represent the estimated *R*_0_, *μ* and *d*

### **Inference of *R*_*0*_, *μ* and *d* by Approximate Bayesian Computation (ABC).**

Due to the complexity of the parameter space, we used Approximate Bayesian Computation (ABC) method by comparing the simulated 10^6^ cell lines with observed data with HeLa to estimate *R*_0_, *μ* and *d*. ABC algorithms are a group of methods for performing Bayesian inference without the need for explicit evaluation of the model likelihood function. The algorithms can be used with implicit computer models that generate sample data sets rather than likelihoods (36, 37). By using ABC we can compute the posterior probability distribution of a multivariate parameter *Θ* (*Θ* ={*R*_0_, *μ*, *d*}). A parameter value *Θ*_*i*_ is sampled from its prior *f*(*Θ*) distribution *f*(*Θ*) to simulate a dataset *S*(*Θ*_*i*_), for i = 1,…, 10^6^. A set of summary statistics, the value that calculated from the data to represent the maximum amount of information in the simplest possible form, *S*(*Θ*_*i*_) is computed from the simulation. By using a distance function *ρ*, we calculated the distance between *S*(*Θ*_*i*_) and observed data *S*_*obs*_. If the distance between *S*(*Θ*_*i*_) and *S*_*obs*_ is less than a given threshold, the parameter value *Θ*_*i*_ is accepted. In order to set the threshold that which simulations are accepted, we provide the tolerance rate ϵ, which is defined as the percentage of accepted simulation (38, 39).

The ABC inference scheme is:

1. Sample a candidate parameter *Θ*_*i*_ ={*R*_0_, *μ*, *d*} from the prior distribution *f*(*Θ*);
2. Simulate the growth process of the cell line *Θ*_*i*_ and calculate the summary statistics; *S*_*i*_
3. Compare the simulated dataset *S*(*Θ*_*i*_), with the observed data *S*_*obs*_, using a distance *p* function and tolerance rate *ϵ*, if *ρ*(*S*(*Θ*_*i*_) *S*_*obs*_) < *ϵ*, accept *Θ*_*i*_;
4. Go to 1.

Here the summary statistics *S*(*Θ*_*i*_)= {simulated cell number of 7 days}. The observed summary statistics *S*_*obs*_={observed 7 days’ cell number from experiment}. The prior distribution *f*(*Θ*) in our model was *R*_0_ ∼ *Uniform* [1.0,3.5], *μ*∼*Uniform* [0.05,0.55] and ∼Uniform [0.01,0.40] The distance function *ρ* we set was the Euclidean distance and the tolerance rate ϵ we set was 0.1%. As a whole 1,000 groups of *Θ*_*i*_ were accepted.

By using different clones from B8 as *S*_*obs*_, we finally calculated the parametric ranges of rapid proliferated cells were *R*_0*f*_ ϵ [2.22, 2.52], *μ*_*f*_ ϵ [0.26, 0.30], and *d*_*f*_ ϵ [0.17,0.20]. Meanwhile, the parametric ranges of defected cells were *R*_0*d*_ ϵ [1.67,2.22], *μ*_*d*_ ϵ[0.28,0.31] and *d*_*d*_ ϵ.[0.17,0.20].

## Acknowledgements

We thank Zhenzhen Liu for the comments and suggestions during the preparation of the manuscript. We thank Xin Wu and Zhipeng Wu for their helps in the artwork of figures.We thank Fang Yang, Chungyan Li and Encheng Dong for their helps in experiments and other support. This study was supported by the Natural Science Foundation of China (91531305 to X.L.), the Strategic Priority Research Program of the Chinese Academy of Sciences (XDB13040300 to X.L. and C.-I.W.), Ministry of Science and Technology, Taiwan (105-2918-I-002 -014 - and 105-2628-B-002 -015 -MY3 to H.Y.W.), Natural Science Foundation grant (31671370, 31301093, 11201224 and 11301294 to T.Z.), the Youth Innovation Promotion Association of Chinese Academy of Sciences (2201080 to T.Z.), National Key Basic Research Program (973 Program) of China (2014CB542006 to C.-I.W.), the Key Research Program of the Chinese Academy of Sciences (KJZD-EW-L14 to X.L.).

## Supporting information

**Fig S1.**
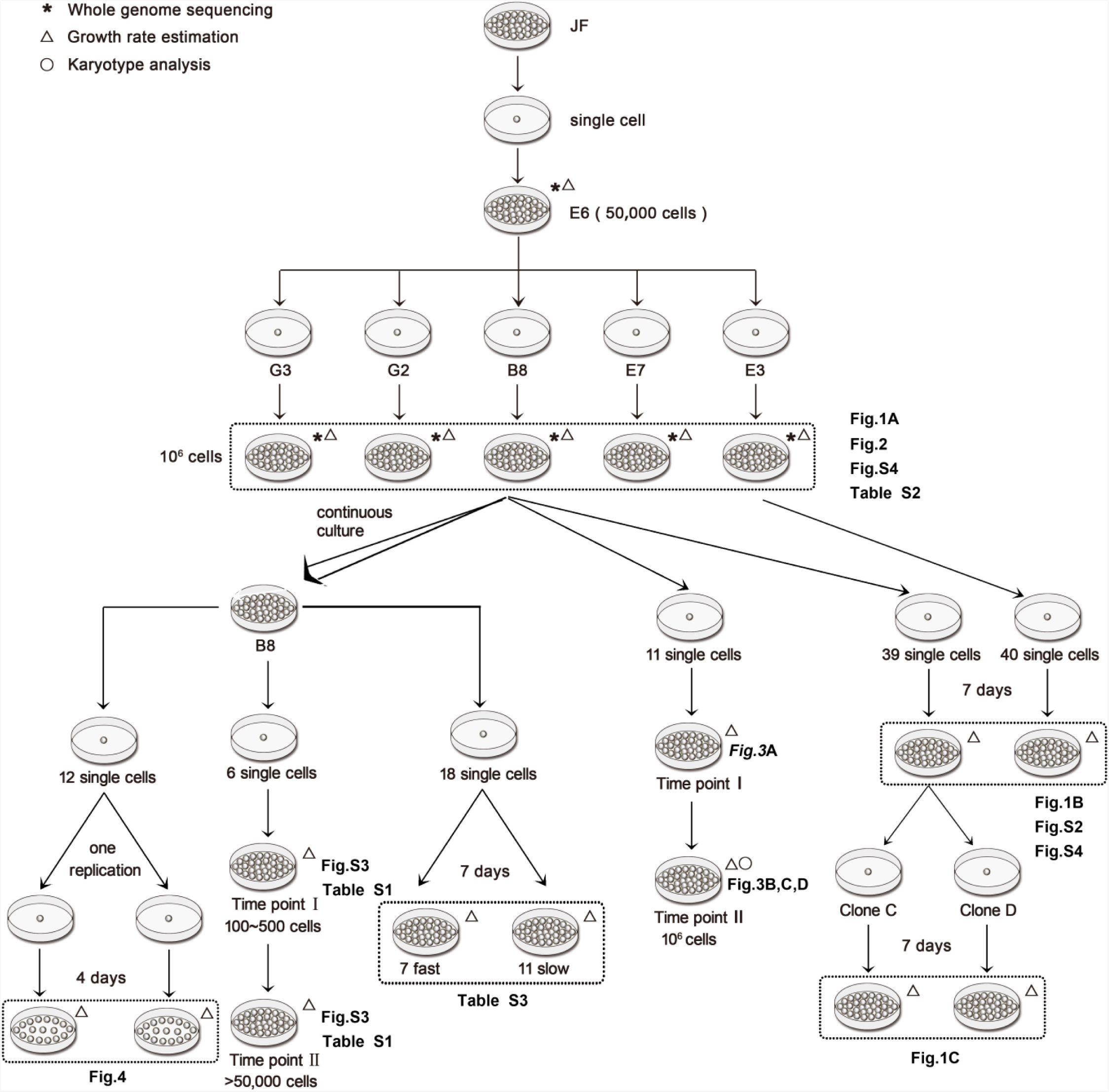
Cell culture and experimental scheme. A single cell (E6) was randomly drew from a Hela cell line (JF) and defined as the ancestral clone. Five clones (B8, G3, E3, E7, G2) were random chosen from E6 when the population size of E6 reached up to > 50,000 cells. The clones B8 and E3 kept cultured for further experiments which are shown in the schematic diagram and described in the Methods and Results.

**Fig S2.**
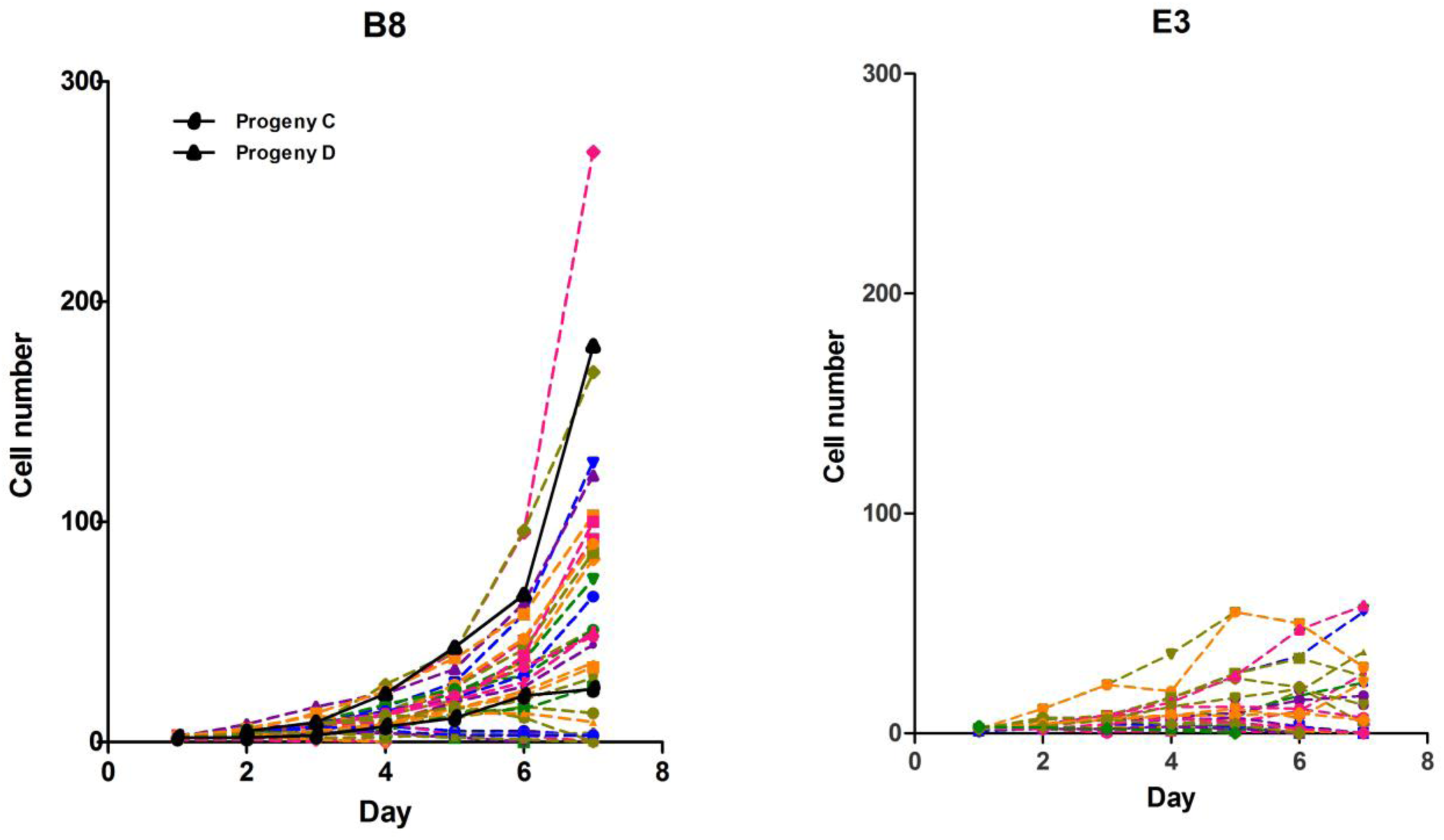
Growth curves of cell clones from B8 and E3. Each curve represents a single cell clone from day 1 to day 7.

**Fig S3.**
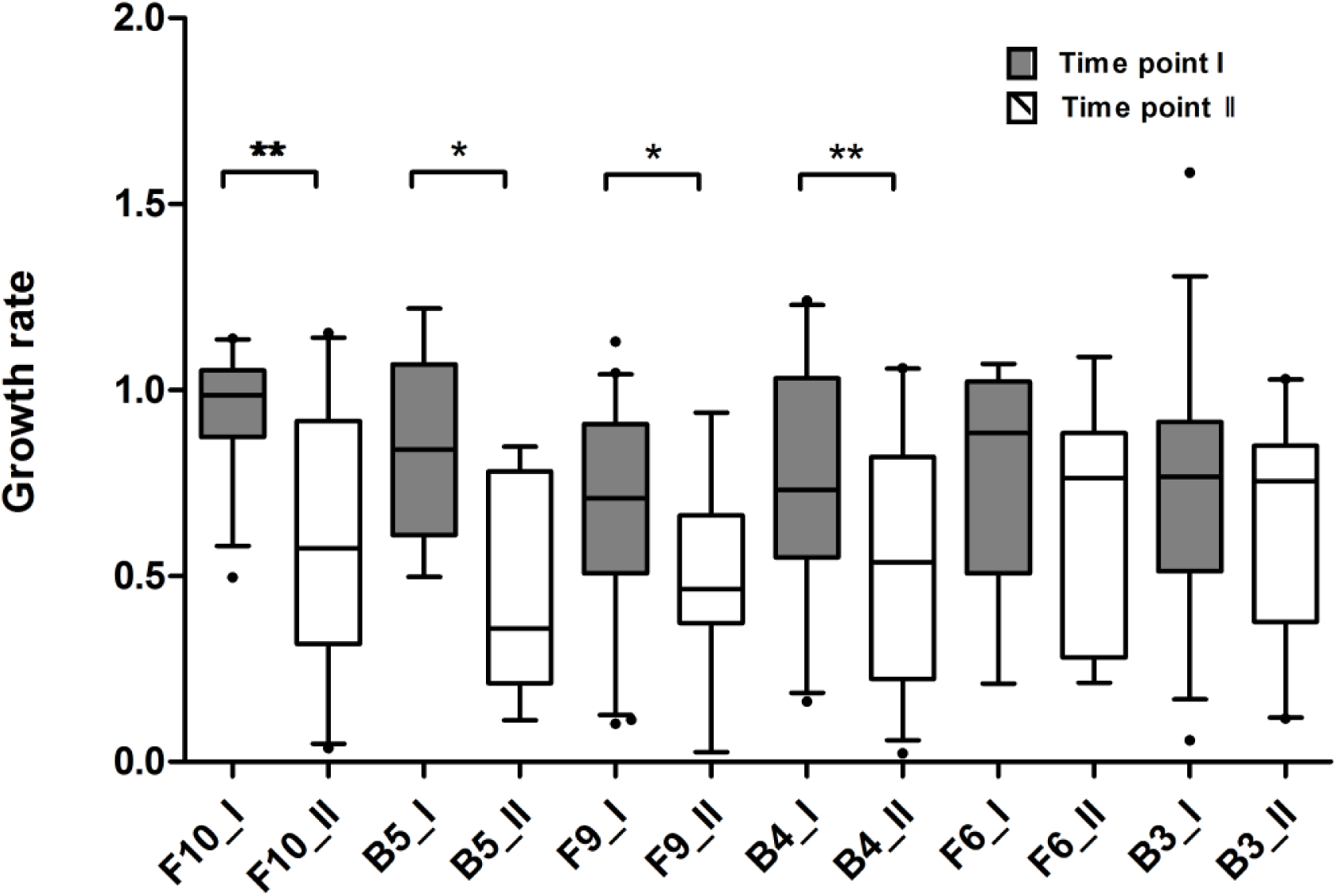
The growth rates of six clones (F10 to B3, labeled on the x-axis) from B8 at the time point I and II. When cell numbers of these six single cell clones reached to 100-500 (after approximately 8-10 cell divisions; Table S1), 20∼30% of cells from each clone were randomly drawn and separated into single cells. The growth of these cells was monitored for 8 days (time point I, solid boxplots). When cell numbers reached to more than 5000 cells (time point II, blank boxplots), a number of single cell clones (Table S1) were generated, and the cell growth for those clones was monitored again. ^*^: p < 0.05; ^**^: p < 0.01, Wilcoxon test.

**Fig S4.**
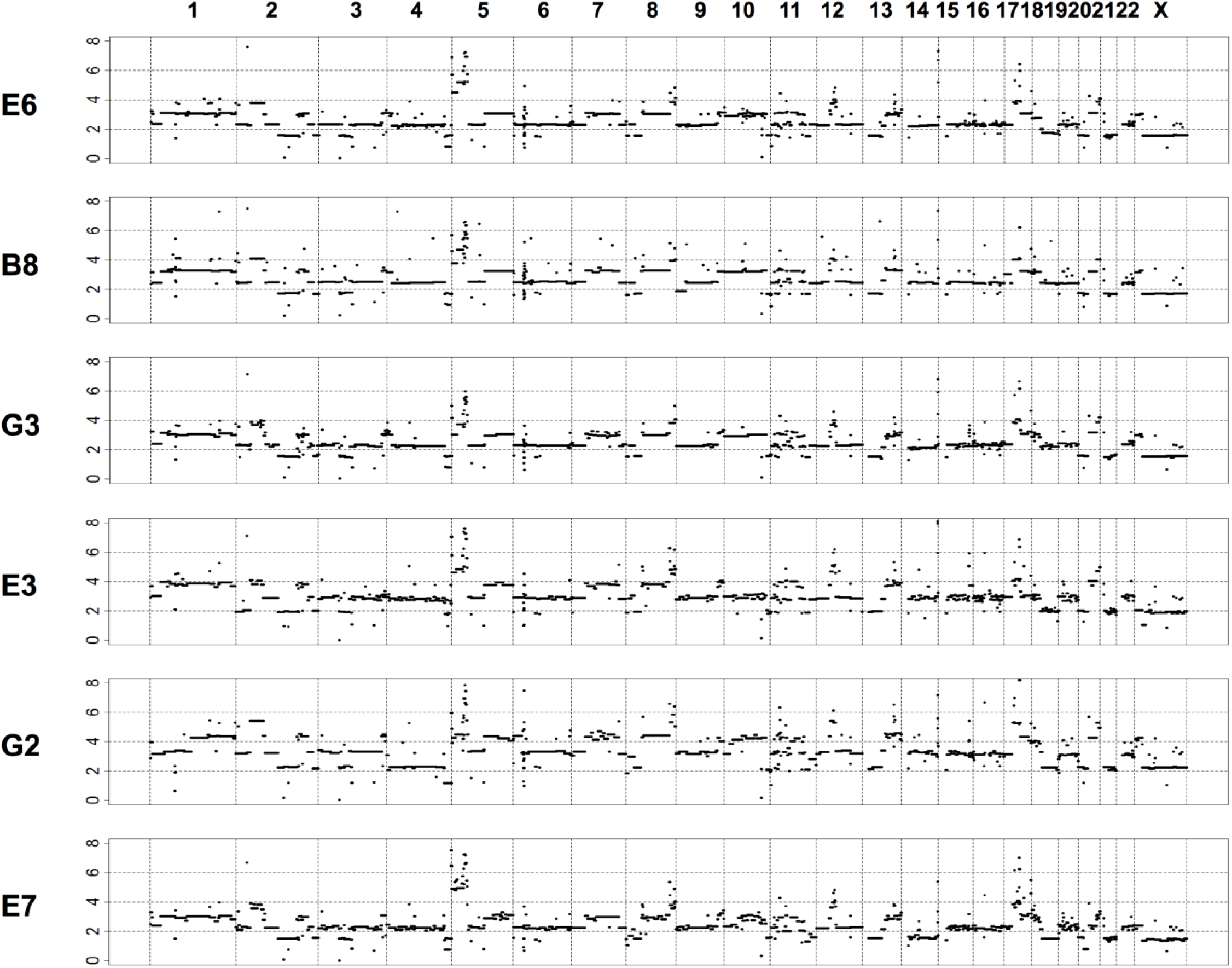
Whole-genome copy number variations in six single cell clones. E6 was originated from a single cell derived from a Hela cell line, JF. G3, B8, G2, E3, E7 were the single cell clones from E6. The copy numbers are shown on the y-axis.

**Fig S5.**
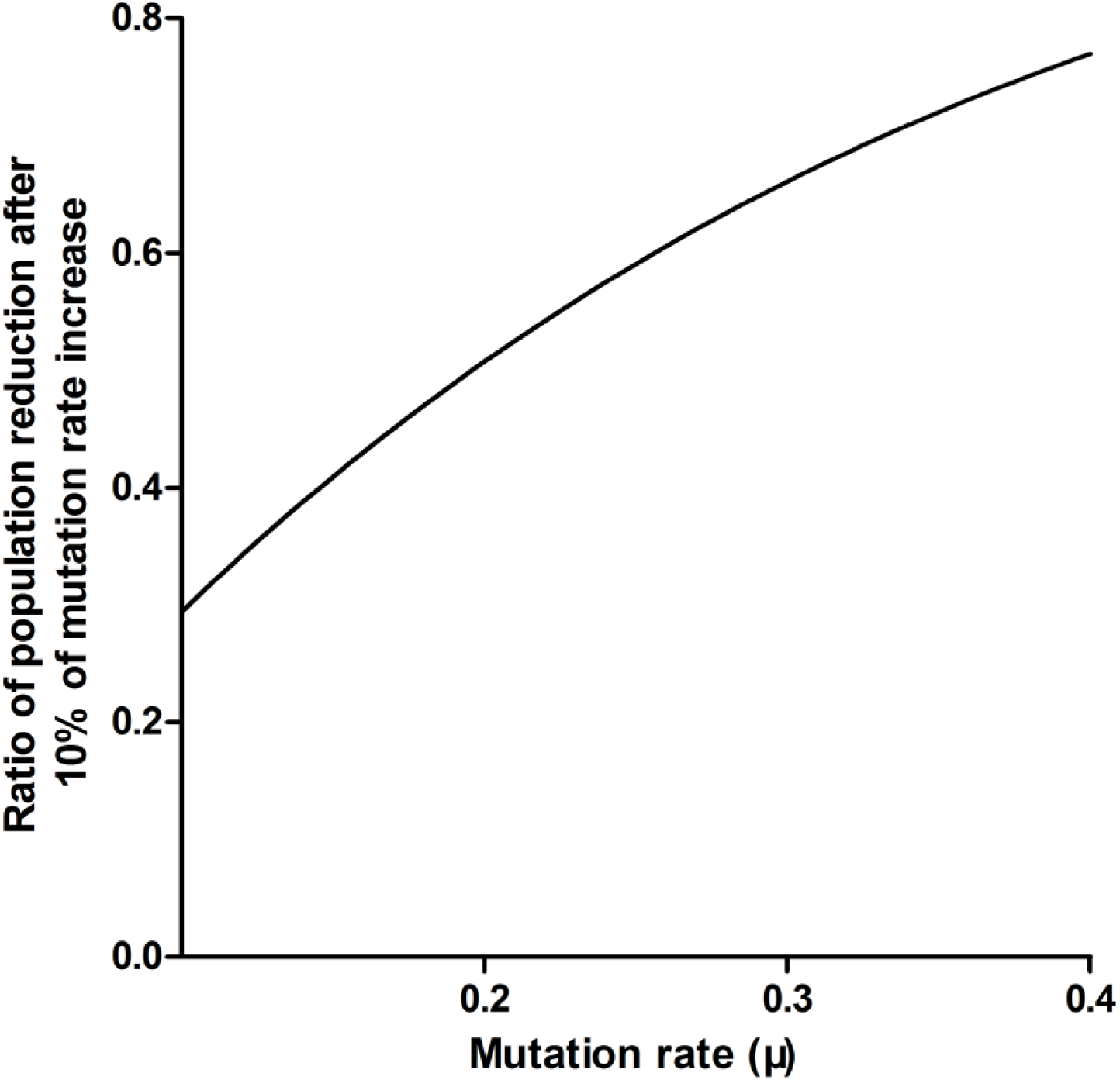
The proportion of cell number reduction with 10% of mutation rate increase. In this figure, the fixed *R*_*0*_ (2.40) and *d* (0.18) are given. The cell number (*N*_*t*_) after 20 days of growing (*t* = 20) with different mutation rate *μ* and 1.1*μ* (*μ* ϵ [0.1,0.4]) were calculated, separately. The y-axes is the reduction ratio of cell numbers derived from 1.1 *μ* and *μ*.

**Table S1.**
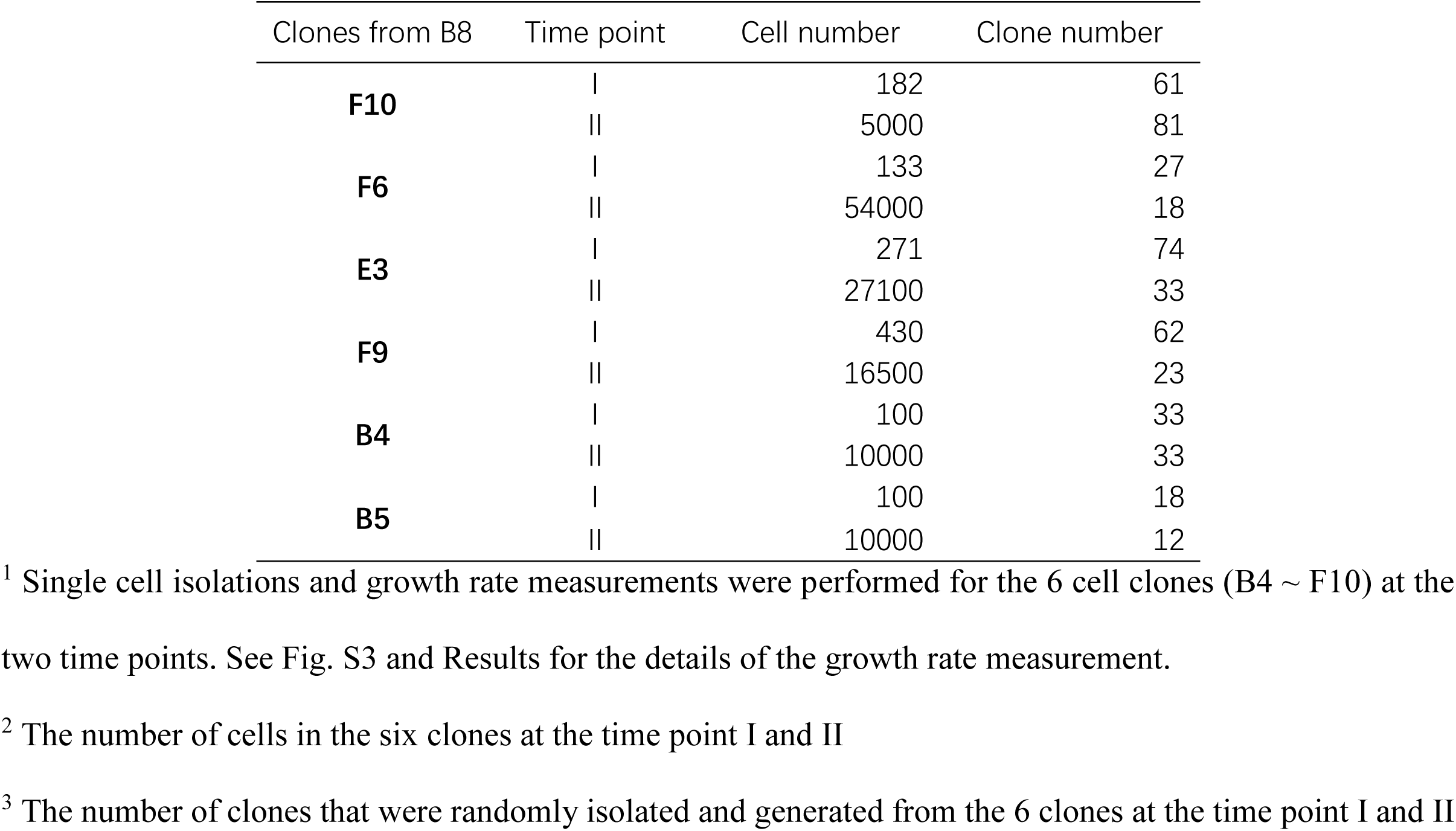
Number of clones that were generated from 6 clones at time point I and II.

**Table S2.** Copy number gain and loss of five descendant clones compared to their ancestor clone (E6).

| Descendant clones from E6 | Chromosome | Start | End | Length | Copy number gain/ loss |
| --- | --- | --- | --- | --- | --- |
| B8 | 2 | 140900000 | 196750000 | 55850000 | gain |
|  | 5 | 0 | 45250000 | 45250000 | loss |
|  | 9 | 200000 | 27500000 | 27300000 | loss |
|  | 14 | 26850000 | 49250000 | 22400000 | gain |
|  | 17 | 0 | 18000000 | 18000000 | gain |
|  | 18 | 0 | 70850000 | 70850000 | gain |
|  | X | 80050001 | 99050000 | 18999999 | gain |
| G3 | 5 | 0 | 47000000 | 47000000 | loss |
|  | 18 | 1700000 | 78077248 | 76377248 | gain |
| E3 | 2 | 0 | 86750000 | 86750000 | loss |
|  | 4 | 0 | 16700000 | 16700000 | loss |
|  | 5 | 0 | 46400000 | 46400000 | loss |
|  | 10 | 0 | 52900000 | 52900000 | loss |
|  | 10 | 53100000 | 123850000 | 70750000 | loss |
|  | 15 | 90450000 | 102531392 | 12081392 | gain |
|  | 16 | 34200000 | 50750000 | 16550000 | gain |
|  | 17 | 22150000 | 57850000 | 35700000 | loss |
|  | 17 | 57950000 | 81195210 | 23245210 | loss |
|  | 18 | 0 | 30900000 | 30900000 | loss |
|  | X | 0 | 33950000 | 33950000 | loss |
| G2 | 1 | 30550000 | 119600000 | 89050000 | loss |
|  | 4 | 0 | 178750000 | 178750000 | loss |
|  | 5 | 0 | 46400000 | 46400000 | loss |
|  | 6 | 0 | 25350000 | 25350000 | gain |
|  | 8 | 0 | 13050000 | 13050000 | loss |
|  | 10 | 0 | 17200000 | 17200000 | loss |
|  | 11 | 60450000 | 71550000 | 11100000 | loss |
|  | 11 | 112000000 | 135006516 | 23006516 | loss |
|  | 18 | 51900000 | 62450000 | 10550000 | gain |
| E7 | 4 | 0 | 16700000 | 16700000 | loss |
|  | 8 | 2500000 | 19150000 | 16650000 | loss |
|  | 10 | 0 | 39000000 | 39000000 | loss |
|  | 11 | 103100000 | 135006516 | 31906516 | loss |
|  | 14 | 0 | 50300000 | 50300000 | loss |
|  | 14 | 52550000 | 107349540 | 54799540 | loss |
|  | 18 | 0 | 12900000 | 12900000 | gain |
|  | 18 | 30850000 | 45250000 | 14400000 | loss |
|  | 18 | 48800000 | 78077248 | 29277248 | loss |
|  | 20 | 15000000 | 63025520 | 48025520 | loss |
|  | X | 0 | 24000000 | 24000000 | loss |

**Table S3.** The number of cells in single cell clones1 from B8 and parameter estimation^2^.

|  | Clone | Day |  |  |  |  |  |  | Parameter estimation |  |  |  |
| --- | --- | --- | --- | --- | --- | --- | --- | --- | --- | --- | --- | --- |
| | | 1 | 2 | 3 | 4 | 5 | 6 | 7 | $R_0$ | $\mu$ | d | $\mu \times d$ |
| Fast growing cells | #1 | 2 | 4 | 8 | 15 | 30 | 45 | 101 | 2.22 | 0.26 | 0.17 | 0.04 |
|  | #2 | 2 | 4 | 8 | 20 | 52 | 99 | 182 | 2.48 | 0.29 | 0.18 | 0.05 |
|  | #3 | 2 | 5 | 10 | 23 | 51 | 84 | 150 | 2.41 | 0.30 | 0.19 | 0.06 |
|  | #4 | 2 | 4 | 8 | 20 | 38 | 57 | 107 | 2.29 | 0.30 | 0.18 | 0.05 |
|  | #5 | 2 | 4 | 14 | 32 | 61 | 112 | 190 | 2.52 | 0.29 | 0.20 | 0.06 |
|  | #6 | 2 | 4 | 12 | 20 | 41 | 100 | 187 | 2.44 | 0.26 | 0.17 | 0.04 |
|  | #7 | 2 | 4 | 8 | 20 | 33 | 55 | 104 | 2.26 | 0.30 | 0.17 | 0.05 |
| Slow growing cells | #8 | 2 | 4 | 7 | 18 | 23 | 35 | 66 | 2.13 | 0.31 | 0.18 | 0.06 |
|  | #9 | 2 | 4 | 7 | 17 | 27 | 46 | 88 | 2.19 | 0.27 | 0.17 | 0.05 |
|  | #10 | 2 | 4 | 7 | 13 | 19 | 27 | 46 | 1.99 | 0.27 | 0.17 | 0.05 |
|  | #11 | 2 | 4 | 6 | 10 | 17 | 30 | 61 | 2.04 | 0.25 | 0.16 | 0.04 |
|  | #12 | 2 | 4 | 8 | 15 | 27 | 48 | 82 | 2.17 | 0.27 | 0.19 | 0.05 |
|  | #13 | 2 | 4 | 4 | 5 | 6 | 9 | 11 | 1.66 | 0.32 | 0.18 | 0.06 |
|  | #14 | 1 | 4 | 6 | 9 | 15 | 25 | 34 | 1.96 | 0.29 | 0.20 | 0.06 |
|  | #15 | 2 | 3 | 5 | 7 | 11 | 13 | 20 | 1.76 | 0.28 | 0.20 | 0.06 |
|  | #16 | 2 | 2 | 4 | 8 | 13 | 19 | 23 | 1.89 | 0.31 | 0.20 | 0.06 |
|  | #17 | 2 | 4 | 8 | 11 | 18 | 26 | 32 | 1.97 | 0.33 | 0.18 | 0.06 |
|  | #18 | 2 | 4 | 8 | 15 | 30 | 51 | 88 | 2.21 | 0.27 | 0.18 | 0.05 |

<sup>1</sup> Single cells were randomly drawn from B8, followed by cell culture for seven days. The cell numbers in the single cell clones were counted every day. According to their cell numbers on the 7<sup>th</sup> day, newly derived cell clones are grouped into fast (cell number > 100) and slow (cell number < 100) growing clones.
<sup>2</sup> The Approximate Bayesian Computation (ABC) method was performed to estimate $R_0$ , $\mu$ and $d$ for each clone.

**Table S4.** R0 estimation with constant μ and d.

|  | Clone | Observed data |  |  |  |  |  |  | R<sub>0</sub> | Expected data |  |  |  |  |  |  | Chi-square test<br>(P-value) |
| --- | --- | --- | --- | --- | --- | --- | --- | --- | --- | --- | --- | --- | --- | --- | --- | --- | --- |
|  |  | Day |  |  |  |  |  |  |  | Day |  |  |  |  |  |  |  |
|  |  | 1 | 2 | 3 | 4 | 5 | 6 | 7 |  | 1 | 2 | 3 | 4 | 5 | 6 | 7 |  |
| B8 | A4 | 2 | 7 | 7 | 16 | 34 | 39 | 70 | 2.13 | 2.1 | 4.3 | 8.2 | 14.9 | 25.6 | 41.8 | 64.5 | 0.672 |
|  | B4-1 | 2 | 4 | 8 | 9 | 10 | 10 | 11 | 1.64 | 1.6 | 2.5 | 3.8 | 5.2 | 6.9 | 8.7 | 10.4 | 0.468 |
|  | B4-2 | 2 | 4 | 8 | 14 | 27 | 44 | 80 | 2.19 | 2.2 | 4.5 | 8.9 | 16.7 | 29.5 | 59.4 | 78.4 | 0.385 |
|  | B6-1 | 4 | 6 | 12 | 20 | 33 | 48 | 78 | 2.18 | 2.2 | 4.5 | 8.8 | 16.4 | 28.8 | 48.0 | 75.9 | 0.773 |
|  | B6-2 |  | 4 | 4 | 11 | 18 | 20 | 69 | 2.16 | 2.2 | 4.4 | 8.6 | 15.8 | 27.5 | 45.4 | 71.2 | 5.208e-08*** |
|  | B9 | 2 | 4 | 8 | 16 | 28 | 50 | 98 | 2.26 | 2.3 | 4.8 | 9.8 | 18.9 | 34.5 | 59.6 | 97.7 | 0.611 |
|  | B10 | 2 | 4 | 7 | 12 | 25 | 27 | 49 | 1.99 | 2.0 | 3.8 | 6.7 | 11.4 | 18.3 | 27.8 | 40.0 | 0.741 |
|  | C10-1 | 2 | 4 | 8 | 16 | 29 | 42 | 72 | 2.17 | 2.2 | 4.5 | 8.7 | 16.0 | 28.1 | 46.7 | 73.5 | 0.994 |
|  | C10-2 | 1 | 2 | 2 | 3 | 5 | 9 | 12 | 1.67 | 1.7 | 2.6 | 4.0 | 5.6 | 7.6 | 9.7 | 11.8 | 0.387 |
|  | D3-1 | 2 | 4 | 6 | 10 | 18 | 36 | 87 | 2.21 | 2.2 | 4.6 | 9.2 | 17.3 | 30.8 | 52.1 | 83.5 | 6.214e-04*** |
|  | D3 -2 | 2 | 3 | 6 | 12 | 24 | 36 | 72 | 2.18 | 2.2 | 4.5 | 8.8 | 16.4 | 28.8 | 48.0 | 75.9 | 0.182 |
|  | D4 | 2 | 4 | 8 | 14 | 18 | 34 | 82 | 2.20 | 2.2 | 4.6 | 9.1 | 17.0 | 30.1 | 50.7 | 80.9 | 0.008** |
|  | D5 | 2 | 4 | 8 | 15 | 20 | 35 | 49 | 2.04 | 2.0 | 3.9 | 7.2 | 12.6 | 20.7 | 32.3 | 47.7 | 0.994 |
|  | D10 | - | 4 | 8 | 12 | 26 | 55 | 125 | 2.33 | 2.3 | 5.1 | 10.8 | 21.4 | 40.2 | 71.6 | 120.9 | 6.323e-04*** |
|  | D11 | 2 | 4 | 8 | 17 | 38 | 54 | 140 | 2.37 | 2.4 | 5.3 | 11.3 | 22.9 | 43.7 | 79.3 | 136.2 | 0.011* |
|  | E3 | - | 4 | 7 | 12 | 24 | 69 | 82 | 2.23 | 2.2 | 4.7 | 9.4 | 17.9 | 32.3 | 55.0 | 89.0 | 0.071 |
|  | E4 | 2 | 6 | 15 | 30 | 50 | 73 | 139 | 2.36 | 2.4 | 5.3 | 11.2 | 22.5 | 42.8 | 77.3 | 132.3 | 0.594 |
|  | E5 | 1 | 4 | 3 | 8 | 17 | 23 | 39 | 1.97 | 2.0 | 3.7 | 6.5 | 10.9 | 17.4 | 26.2 | 37.4 | 0.352 |
|  | E7 | 2 | 4 | 7 | 9 | 12 | 18 | - | 1.82 | 1.8 | 3.1 | 5.1 | 8.0 | 11.7 | 16.3 | 21.5 | 0.961 |
|  | F2 | 2 | 4 | 7 | 15 | 26 | 51 | 79 | 2.19 | 2.2 | 4.5 | 8.9 | 16.7 | 29.5 | 59.4 | 78.4 | 0.852 |
|  | F3 | 2 | 4 | 8 | 16 | 33 | 64 | 117 | 2.31 | 2.3 | 5.1 | 10.5 | 20.6 | 38.5 | 68.0 | 113.9 | 0.717 |
|  | F7 | 2 | 3 | 6 | 11 | 17 | 26 | 48 | 2.01 | 2.0 | 3.8 | 6.9 | 11.8 | 19.2 | 29.5 | 43.0 | 0.946 |
|  | F7 | 2 | 2 | 3 | 7 | 11 | 21 | 24 | 1.89 | 1.9 | 3.4 | 5.7 | 9.3 | 14.1 | 20.4 | 27.9 | 0.458 |
|  | E8 | 2 | 3 | 6 | 10 | 14 | 14 | 17 | 1.76 | 1.8 | 2.9 | 4.6 | 7.0 | 9.9 | 13.3 | 17.0 | 0.865 |
|  | F10 | - | 5 | 9 | 22 | 43 | 67 | 180 | 2.45 | 2.5 | 5.7 | 12.5 | 26.1 | 51.6 | 96.8 | 171.9 | 3.548e-03** |
|  | G2 | 2 | 7 | 15 | 28 | 53 | 97 | 230 | 2.54 | 2.5 | 6.1 | 14.0 | 30.2 | 61.8 | 120.2 | 221.3 | 0.252 |
|  | G12 | - | 4 | 5 | 5 | 8 | 9 | 16 | 1.68 | 1.7 | 2.7 | 4.0 | 5.8 | 7.8 | 10.1 | 12.3 | 0.883 |
|  | H3 | 1 | 3 | 6 | 8 | 12 | 20 | 34 | 1.92 | 1.9 | 4.5 | 6.0 | 9.9 | 15.3 | 22.4 | 31.2 | 0.752 |
|  | H6 | 2 | 4 | 8 | 12 | 19 | 22 | 40 | 1.94 | 1.9 | 3.6 | 6.2 | 10.3 | 16.1 | 23.9 | 33.5 | 0.884 |
| E3 | C3 | 1 | 2 | 3 | 8 | 16 | 19 | 32 | 1.93 | 1.9 | 3.5 | 6.1 | 10.1 | 15.7 | 23.1 | 32.4 | 0.361 |
|  | C4-1 | 2 | 4 | 9 | 18 | 30 | 46 | 62 | 2.12 | 2.1 | 4.3 | 8.1 | 14.6 | 25.1 | 40.6 | 62.4 | 0.901 |
|  | C4-2 | 1 | 6 | 8 | 16 | 27 | 34 | - | 2.11 | 2.1 | 4.2 | 8.0 | 14.4 | 24.5 | 39.5 | 60.4 | 0.695 |
|  | C6 | 2 | 2 | 4 | 8 | 9 | 17 | 23 | 1.83 | 1.8 | 3.2 | 5.2 | 8.1 | 12.0 | 16.8 | 22.3 | 0.908 |
|  | D3 | 2 | 4 | 8 | 15 | 22 | 41 | 63 | 2.11 | 2.1 | 4.2 | 8.0 | 14.4 | 24.5 | 39.5 | 60.4 | 0.998 |
|  | D7-1 | 2 | 2 | 4 | 7 | 18 | 22 | 49 | 2.04 | 2.0 | 3.9 | 7.2 | 12.6 | 20.7 | 32.3 | 47.7 | 0.028* |
|  | D7-2 | - | 4 | 8 | 14 | 26 | 47 | 58 | 2.10 | 2.1 | 4.2 | 7.9 | 14.1 | 23.9 | 38.4 | 58.4 | 0.882 |
|  | D8 | 2 | 4 | 8 | 13 |  | 20 | 42 | 1.96 | 2.0 | 3.6 | 6.4 | 10.7 | 16.9 | 25.4 | 36.0 | 0.687 |
|  | E9 | 2 | 4 | 4 | 8 | 14 | 18 | 18 | 1.78 | 1.8 | 3.0 | 4.8 | 7.3 | 1.1 | 14.2 | 18.4 | 0.039* |
|  | E10 | 2 | 4 | 7 | 7 | 9 | 15 | 17 | 1.74 | 1.7 | 2.9 | 4.5 | 6.6 | 9.3 | 12.4 | 15.7 | 0.935 |
|  | F4 | 1 | 2 | 2 | 4 | 7 | 19 | 37 | 1.97 | 2.0 | 3.7 | 6.5 | 10.9 | 17.4 | 26.2 | 37.4 | 1.364e-07*** |
|  | F7 | - | 4 | 6 | 8 | 10 | 16 | 21 | 1.77 | 1.8 | 3.0 | 4.7 | 7.1 | 10.1 | 13.8 | 17.7 | 0.918 |
|  | F9 | 2 | 4 | 8 | 16 | 27 | 35 | 55 | 2.06 | 2.1 | 4.0 | 7.4 | 13.1 | 21.7 | 34.2 | 51.1 | 0.928 |
|  | G3 | 2 | 4 | 7 | 16 | 32 | 48 | 86 | 2.21 | 2.2 | 4.6 | 9.2 | 17.3 | 30.8 | 52.1 | 83.5 | 0.967 |

Note: Using ABC method with the fixed $\mu$ (0.29) and $d$ (0.18), $R_0$ was estimated from the cell clones derived from B8 and E3. The expected data were calculated from equation (1) using the estimated $R_0$ . Chi-square test was used to test whether the expected data are fitted with the observed data. \*, P-value < 0.05; \*\*, P-value < 0.01; \*\*\*, P-value < 0.001.

